# Glycosaminoglycans Modulate Long-Range Mechanical Communication Between Cells in Collagen Networks

**DOI:** 10.1101/2021.09.13.460122

**Authors:** Xingyu Chen, Dongning Chen, Ehsan Ban, Paul A. Janmey, Rebecca G. Wells, Vivek B. Shenoy

## Abstract

Cells can sense and respond to mechanical forces in fibrous extracellular matrices (ECM) over distances much greater than their size. This phenomenon, termed long-range force transmission, is enabled by the realignment (buckling) of collagen fibers along directions where the forces are tensile (compressive). However, whether other key structural components of the ECM, in particular glycosaminoglycans (GAGs), can affect the efficiency of cellular force transmission remains unclear. Here we developed a theoretical model of force transmission in collagen networks with interpenetrating GAGs, capturing the competition between tension-driven collagen-fiber alignment and the swelling pressure induced by GAGs. Using this model, we show that the swelling pressure provided by GAGs increases the stiffness of the collagen network by stretching the fibers in an isotropic manner. We found that the GAG-induced swelling pressure can help collagen fibers resist buckling as the cells exert contractile forces. This mechanism impedes the alignment of collagen fibers and decreases long-range cellular mechanical communication. We experimentally validated the theoretical predictions by comparing collagen fiber alignment between cellular spheroids cultured on collagen gels versus collagen-GAG co-gels. We found significantly less alignment of collagen in collagen-GAG co-gels, consistent with the prediction that GAGs can prevent collagen fiber alignment. The roles of GAGs in modulating force transmission uncovered in this work can be extended to understand pathological processes such as the formation of fibrotic scars and cancer metastasis, where cells communicate in the presence of abnormally high concentrations of GAGs.

**Statement of significance:** Glycosaminoglycans (GAGs) are carbohydrates that are expressed ubiquitously in the human body and are among the key macromolecules that influence development, homeostasis, and pathology of native tissues. Abnormal accumulation of GAGs has been observed in metabolic disorders, solid tumors, and fibrotic tissues. Here we theoretically and experimentally show that tissue swelling caused by the highly polar nature of GAGs significantly affects the mechanical interactions between resident cells by altering the organization and alignment of the collagenous extracellular matrix. The roles of GAGs in modulating cellular force transmission revealed here can guide the design of biomaterial scaffolds in regenerative medicine and provides insights on the role of cell-cell communication in tumor progression and fibrosis.

## Introduction

The extracellular matrix (ECM) in tissues and organs consists of collagen, enzymes, glycoproteins, and other macromolecules that provide structural and biochemical support to the cellular constituents. Structurally, the ECM is principally composed of an interlocking meshwork of collagen and glycosaminoglycans^1^ (GAGs), which are large linear polysaccharides consisting of repeating disaccharide units. Collagen forms a crosslinked network of semi-flexible fibers. A key feature of the fibrous nature of the collagen network is that cells can transmit and sense forces over distances much larger than their size^2^. This long-range transmission of forces is necessary for various physiological and pathological processes. For example, cancer cells generate sufficiently high force to align nearby ECM fibers, which promote cell migration and diffusion of growth factors from the tumor microenvironment^3^. In general, collagen fibers reorient and align along the directions where the strains are tensile and buckle in the directions in which the strains are compressive. This mechanism allows the network to stiffen in an anisotropic manner along the direction of tensile principal strain as the network is deformed^2^. The aligned fibers provide efficient paths for force transmission. The alignment of ECM fibers further increases cancer cell and ECM stiffness and, therefore, the traction forces that the cells exert on the surrounding ECM create a positive feedback loop by upregulating mechano-sensitive signaling, which further enhances the ability of the cells to exert forces^3^. While previous studies have attempted to model the mechanical behavior and force transmission in tissues by considering the effect of GAGs^4,5^ interpenetrating the collagen network, the impact of GAGs on cellular force transmission remains unresolved.

Elevated levels of GAGs, specifically hyaluronic acid (HA), are observed in many types of tumors, wound sites, and other pathological processes such as fibrosis. In grade 3 human ovarian tumors, for example, median HA concentrations are 0.197 mg/mL (with a maximum of 4 mg/mL), over 50 times higher than in benign tumors (0.003mg/mL)^6^. Excess HA accumulation has also been reported in human lung cancer, with a 4- to 200-fold increase^6^, and in human prostate cancer, with a 7-fold increase^6^. The distribution of HA varies significantly from point to point in the tumor microenvironment, and high concentrations of HA are associated with cancer cell invasiveness. In breast tumors, enrichment of HA has been observed exclusively at the leading edge of the tumor^7^. The accumulation and localization of HA in stroma are linked to reduced survival^7^, suggesting that HA participates in the generation of the pro-cancerous stroma. A quantitative analysis of cellular force transmission in tissues with GAGs thus has the potential to provide insights into cellular mechanosensing in pathological microenvironments.

In this study, we developed a model that captures both the non-linear strain stiffening behavior of the collagen network and the swelling induced by GAGs. We used this model to analytically and numerically predict the range of cellular force transmission in fibrous matrices that contain different concentrations of GAGs. Our model predicts that GAGs can significantly modulate long-range force transmission and mechanical communication between cells in collagen networks by making it difficult for fibers to align in an anisotropic manner. We quantitatively validated the theoretical predictions by examining the alignment of collagen fibers between cell spheroids cultured on collagen gels and collagen-HA co-gels. Our work fills an important gap in understanding cellular mechano-sensing by shedding light on hitherto unexplored mechanisms of cellular communication in pathological conditions that involve GAG overproduction, including cancer, fibrosis, and wound healing.

## Results

### A chemo-mechanical model of cellular force transmission in collagen networks with interpenetrating GAGs

We considered the ECM as a mixture of three components: a solid component (collagen network with interpenetrating GAGs), an incompressible fluid component (interstitial fluid), and ions (cations and anions). GAGs contain a large number of acidic groups (e.g., the carboxyl group on HA), which can dissociate into mobile protons and fixed negative charges^8^. Each fixed negative charge requires a counter-ion nearby to maintain charge-neutrality. In physiological conditions, the total ion concentration of the ECM is always greater than that of the external ionic solution^9^. *In vivo*, external ionic strength is maintained by the circulatory system, including blood vessels and lymphatics. For example, sodium concentration is maintained at around 150 mmol/mL in blood^10^. The difference between the ion concentration in the ECM and the surrounding environment gives rise to the well-known Donnan osmotic pressure^11^, which drives water flux into the ECM. Since GAGs are large, charged molecules that are physically restrained in the dense collagen network, the influx of water can cause the ECM to swell. In the physiological state of 150 mmol/mL NaCl, the ECM is swollen, with a swelling pressure resisted by the elastic stress in the collagen-GAG solid matrix. Depending on the volume fraction of GAGs in the ECM and the concentration of ions in the surrounding environment, the matrix can swell to different levels as GAGs attract water into the matrix (Fig. 1a). For example, Lai et al.^5^ showed that collagen-HA co-gels were swollen in hypo-osmotic solutions while collagen gels were not; the swelling also decreased with increasing ion concentrations in the external solution. Voutouri et al.^12^ identified a linear correlation between tumor swelling and the ratio of HA to collagen content. To predict the impact of GAG-induced swelling on cellular force transmission, we developed a constitutive model for collagen-GAG co-gels described by

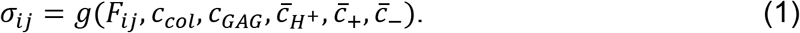

**Figure 1.**
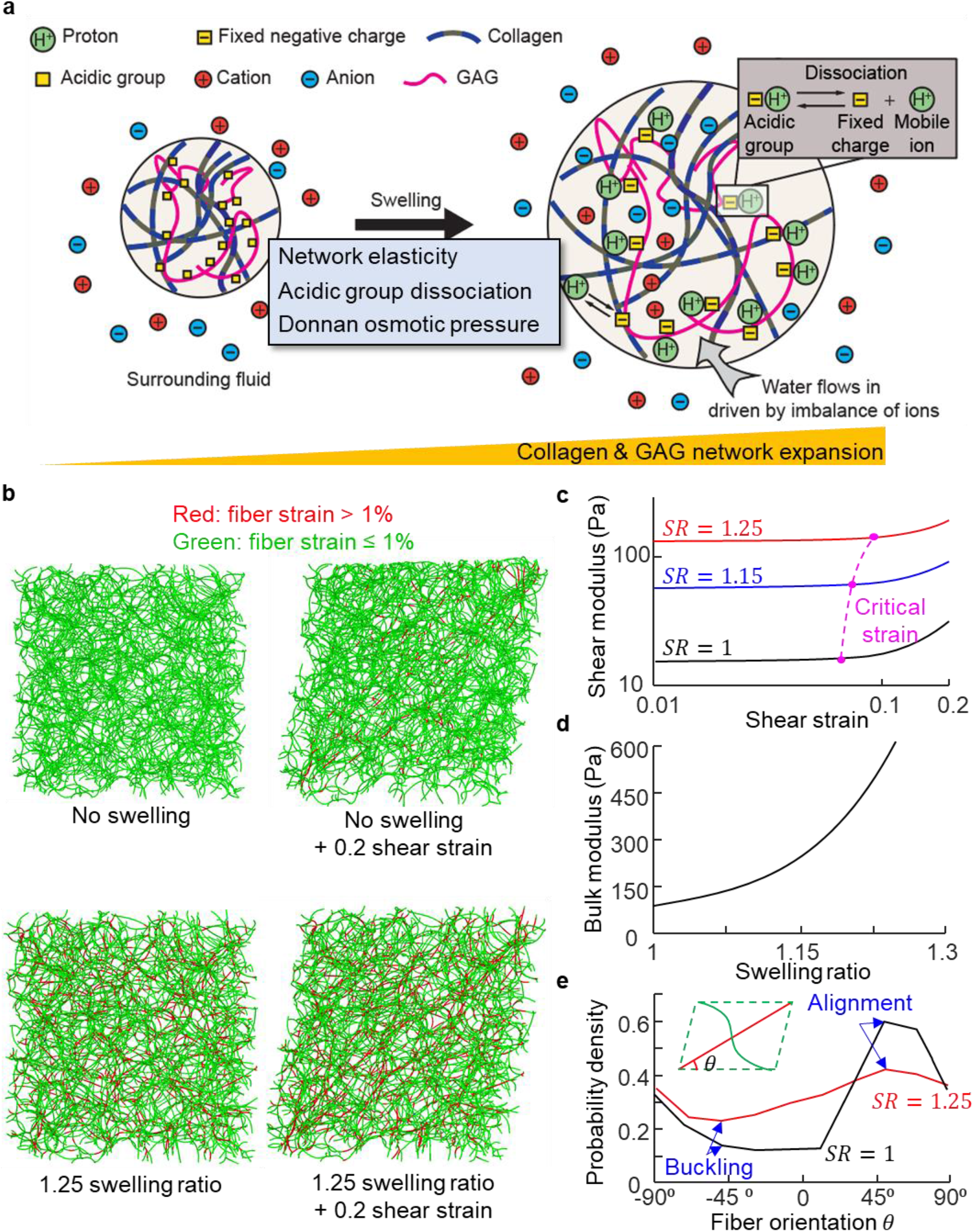
(a) Schematic showing free swelling of a collagen-GAG gel. The carboxyl groups dissociate into negative charges that are fixed to the GAG backbone and protons in solution, which increases the osmolarity in the gel. Ions and water from the surrounding solvent flow into the gel (when the system reaches steady state). (b) 3D discrete fiber simulations of random fiber networks before (left) and after (right) shear deformation (20% shear strain). The networks in the bottom row are swollen (25% volumetric strain) prior to the application of shear deformation. (c) Shear modulus as a function of the shear strain for networks with initial swelling ratios (SR) of 1, 1.15, and 1.25, obtained from discrete fiber simulations. (d) The bulk modulus of the collagen network as a function of the swelling ratio. (e) Distribution of orientation of stretched fibers (strain > 1%) after 20% shear for networks with SR = 1 and 1.25. The high (low) probability of stretched fibers along 45° (−45°) directions indicates fiber alignment (buckling).

In this equation, *σ_ij_* and *F_ij_* denote the stress tensor and the deformation gradient of the collagen-GAG network, respectively. The parameters characterizing the co-gels include concentrations of collagen (*c_col_*) and GAGs (*c_GAG_*) in the ECM and the concentrations of hydrogen ions 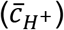, cations 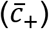, and anions 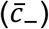 in the surrounding environment.

To obtain the exact form of *g*() in Eq. 1, we considered three major mechanisms involved in the swelling process: 1) acidic groups on GAGs dissociate into fixed negative charges and mobile protons, increasing the ionic concentration in the matrix, 2) water flow into the matrix driven by Donnan osmotic pressure to compensate for the increased ion concentration, and 3) the network of collagen and GAGs is stretched as water flows in. To capture these mechanisms, we first incorporated the Donnan osmotic pressure induced by fixed negative charges, and then introduced the constitutive relation of the collagen-GAG network that is stretched due to swelling. Using these constitutive relations, we simulated cell contraction in the co-gels and studied the impact of GAGs on cellular force transmission.

#### 1. Swelling pressure induced by the GAGs

GAGs contain acidic groups (e. g., carboxyl group) that can dissociate into mobile protons and negative charges (–*COO*^−^) fixed to the GAG chains (Fig. 1a). The dissociation can be calculated by the detailed balance condition that relates the concentrations of the reactants (undissociated acid groups) to the products (protons and carboxyl groups)^13^,

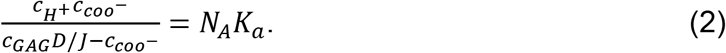

Here *c_C00^−^_* = *c_H^+^_* + *c*_+_ – *c*_−_ denotes the concentration of fixed charges (assuming electrical neutrality), *c_H^+^_, c*_+_ and *c*_−_ denote the concentration of protons, cations, anions in the matrix, respectively, *N_A_* denotes the Avogadro number, and the dissociation constant *K_a_* is a measure of the strength of the acidic group; *D* denotes the number of acidic groups per unit mole of GAGs and *J* = det (*F_ij_*) denotes the ratio between the volumes of the gel after and before deformation. The fixed negative charges can cause an increase in the concentration of mobile positive ions in the matrix and attract water from the surrounding environment. The driving force for inflow of water due to the imbalance of ion concentrations is the osmotic pressure, Π, that can be calculated using the relation^11^:

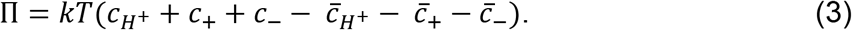

Here 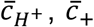 and 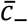 denote the concentrations of protons, cations, and anions in the surrounding environment, respectively. The concentrations of ions within the matrix and in the external solution are related by the Donnan equations^11^, which can be obtained by balancing the chemical potentials of the mobile ionic species in the ECM and in the external solution,

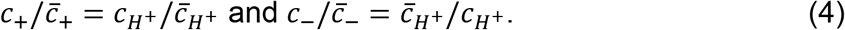

Solving Eq. 2–4 yields the expression of the swelling pressure induced by the acidic groups on GAGs^13^,

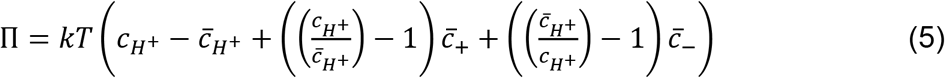

where,

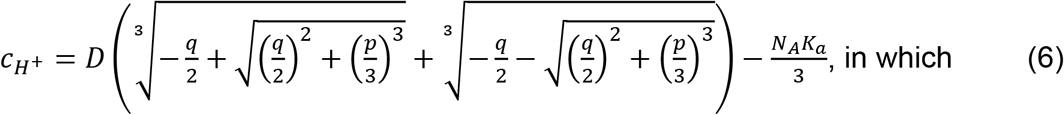

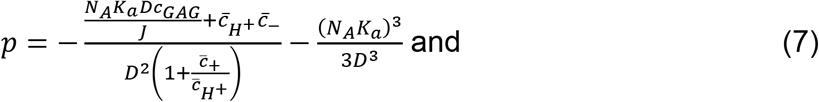

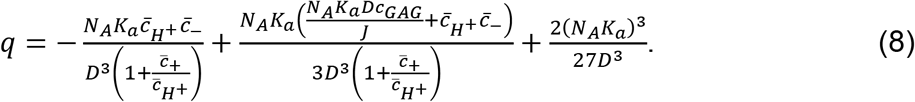

The above equation relates the osmotic pressure in the matrix to the ionic concentration of the external solution, the density and strength of acidic groups on the GAGs and the overall swelling ratio (J), which in turn depends on the mechanics of the GAG-collagen network, which we consider next.

#### 2. The elasticity of the collagen network with interpenetrating GAGs

Both the collagen network and GAGs contribute to the elasticity of the matrix. GAG chains interact with the collagen network through hydrogen bonds and electrostatic interactions^14,15^. Additionally, GAGs can be covalently coupled to a core protein *in vivo* to form proteoglycans^16^. Depending on the type of the core protein, proteoglycans can noncovalently bind to collagen fibrils^15^. Given the interactions between the GAGs and collagen, it is reasonable to assume that GAGs and collagen undergo the same state of deformation on a macroscopic length scale (on the order of cell dimensions (~10-20 μm) or more). We considered the strain energy density of the collagen-GAG network (*W_E_*) as the sum of the strain energy density of the collagen (*W_col_*) and the GAGs (*W_GAG_*):

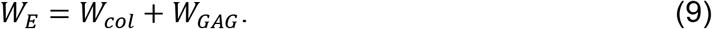

The mechanical behavior of the flexible GAG chains is dominated by their configurational entropy^17^, which can be captured by the non-fibrous (Neo-Hookean) energy density function

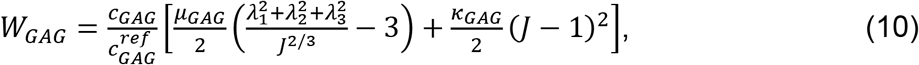

where *λ*_1_, *λ*_2_ and *λ*_3_ denote the stretches in three principal directions and *J* = *λ*_1_*λ*_2_*λ*_3_ denotes the volume change as the matrix deforms. The stiffness of the GAG network depends on the concentration of GAGs; we explicitly scale the strain energy density with the concentration of the GAGs. Since GAGs in physiological conditions are dilute (on the scale of mg/mL), we explore the change of concentration in the physiologically relevant range by adopting a linear scaling relation to estimate the impact of concentration on stiffness. Here, *c_GAG_* denotes the concentration of GAGs in the undeformed configuration, and 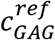 denotes a scaling factor that represents the GAG concentration at which the shear and bulk moduli are *μ_GAG_* and *κ_GAG_*, respectively.

The collagen network was modeled using the constitutive relation we recently developed^2,18,19^, which can account for the fact that collagen fibers can buckle, stretch, and align, depending on the nature of the applied stresses. When a randomly oriented collagen network is subject to a general state of stress, a portion of the fibers reorient and align along directions where the strains are tensile, while the rest of the fibers remain random (Fig. 1b). The alignment of and concomitant stretching of fibers in tension make the network increasingly difficult to deform (Fig. 1c). In orientations where fiber strains are compressive, the fibers buckle and provide no resistance to deformation (Fig. 1e). Due to fiber realignment and buckling, the initially isotropic network stiffens in an anisotropic manner. We capture the strain-stiffening mechanism by writing the strain energy density function as a sum of the contributions from aligned fibers and unaligned fibers,

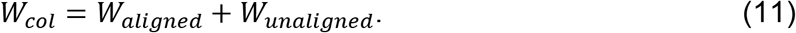

We used the non-fibrous material model (Neo-Hookean) to capture the isotropic mechanical behavior of the randomly aligned fibers and a highly non-linear function 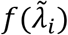 to capture the strain-stiffening effect induced by fiber alignment along tensile principal directions (Fig. S5).

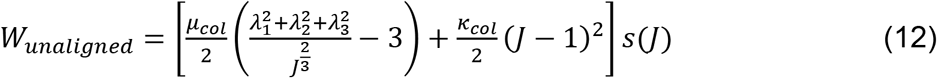

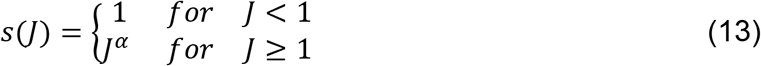

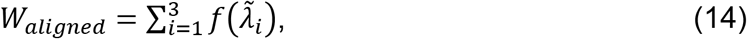

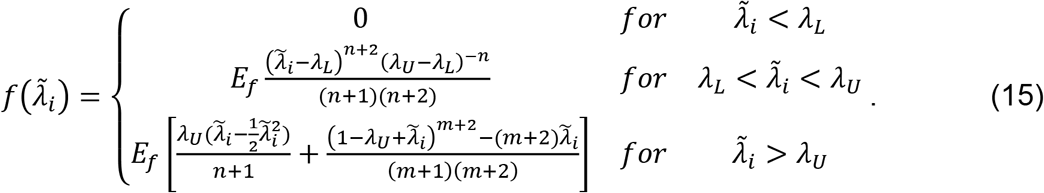

Here, 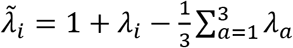 denotes the deviatoric part of the principal stretch. *μ_col_* and *κ_col_* are the shear modulus and bulk modulus of the unaligned “background” fibers, respectively, *E_f_* characterizes the stiffness attributed to the network due to fiber realignment, *m* is the strain-stiffening exponent, *λ_C_* denotes the critical stretch that characterizes the transition to alignment, and *λ_L_* = *λ_C_* – (*λ_C_* – 1)/8 and *λ_U_* = *λ_C_* + (*λ_C_* – 1)/8 denote the lower and upper bounds of the transition range 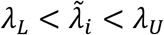, respectively. Once the tensile principal strain goes beyond the transition range, the fibers start to reorient and align with the tensile principal directions, and *n* is an exponent that characterizes the increase in strain energy in the transition range.

When the network experiences volumetric change induced by swelling, the fibers experience tensile forces irrespective of their initial orientation. We found through discrete fiber simulations that the network becomes stiffer in an isotropic manner without fiber alignment. In this case, both the shear modulus (Fig. 1c) and the bulk modulus (Fig. 1d) increased with swelling. To capture this effect, we added an additional scaling factor *s*(*J*) that only depends on the volumetric change given by Eq. 13. The parameter *α* characterizes the non-linear relation between collagen network isotropic stiffness and swelling. Note that, in swollen states, the level of shear or uniaxial loading needed to align fibers is no longer *λ_C_* (using swollen state as the reference configuration, Fig. 1c). The fibers in the network are more difficult to align because fibers are stretched by the GAG-induced swelling, and are more resistant to buckling; thus, a higher level of loading is needed to overcome the tension in the network. To illustrate this point, we compared the orientations of fibers in swollen and unswollen networks using discrete fiber simulations (Fig. 1e). Under the same level of shear strain, a lower percentage of the fibers aligned along the 45° orientation (direction of maximum principal stretch) for the swollen network. The distribution of fiber orientation is flat, indicating that the anisotropy of stiffness decreases with volumetric expansion. As a result, the critical strain needed for fiber alignment increases with the network volumetric expansion (Fig. 1c). Since in this study we focused on moderate levels of swelling (~15% swelling) where the critical strain varies slightly, we kept *λ_c_* constant and used the criterion 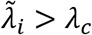 to determine whether the fibers became aligned at a given material point. Note that the parameters *μ_col_, κ_col_, E_f_* and *λ_c_* in Eq. 12–14 depend on the concentration of collagen, *c_col_*. For example, previous studies have identified that the pore size of a collagen network, which determines the critical strain *ϵ_c_*, is inversely proportional to the square root of the collagen concentration^20,21^. In this study, we conducted experiments to calibrate the model specifically for a fixed collagen concentration *c_col_* = 2.5 *mg/mL*.

With the strain energy density of collagen and GAGs, the stress that arises from the elasticity of the matrix 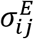 can be calculated as

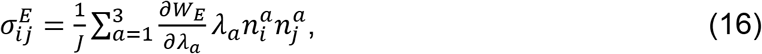

where *λ_a_*(*a* = 1,2,3) and 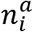 denotes the stretches and the unit vectors along the three principal directions, respectively. Combining the swelling pressure (Eq. 5) with the elasticity of the matrix (Eq. 15), we obtained the overall stress (*σ_ij_*) in the ECM, which can be written as

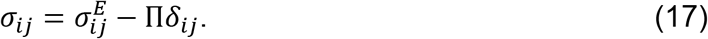

Having developed the model, we first used this constitutive model to predict the dependence of the free swelling behavior on the GAG concentration. Next, by comparing the swelling ratio and shear stiffness of the collagen-GAG network with experiments, we determined the parameters in our constitutive model.

### Collagen matrix swells due to osmotic pressure induced by GAGs

We started analyzing the impact of GAGs by considering the case of free swelling where the ECM was subject to no constraining forces. Since ion concentrations are maintained at a relatively constant level in physiological conditions, we set 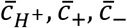 on the outer boundary to be 0.0001 mmol/L (pH = 7), 150 mmol/L, 150 mmol/L, respectively. Enforcing the conditions of mechanical equilibrium,

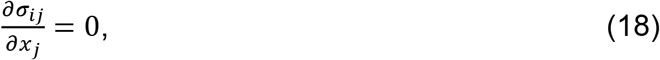

led to *σ_ij_* = 0 throughout the gel^13^, and 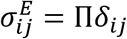 (Eq. 17), where *x_j_* (*j* = 1,2,3) denote the spatial coordinates. This result suggests that the swelling pressure Π induced by GAGs leads to isotropic tensile stress on the solid matrix, leading to elastic deformation of the collagen-GAG network. Since the swelling pressure Π (Eq. 5) depends on the GAG-induced imbalance of ion concentrations across the network boundary, our model predicted that high concentration of GAGs, *c_GAG_*, can induce high swelling pressure Π, and thus requires a large solid matrix stress 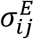 to balance it, causing the expansion of the whole network.

To verify the prediction that GAGs can induce swelling of the collagen network, we conducted free swelling experiments using collagen-HA gels (see materials and methods section). Notably, we used non-crosslinked HA of large molecular size (1.5 MDa). We expect that the HA chains are physically trapped inside the collagen mesh. HA contains carboxyl groups^22^ that can dissociate into fixed negative charges and mobile protons, increasing ion concentration within the collagen-HA gel. The imbalance of ion concentrations inside and outside the gel can drive in water and expand the collagen-HA gel (Fig. 2a). We found that while the collagen gels only showed negligible swelling ratios (defined in the materials and methods section, Eq. 21) of 1.02 ± 0.03, the collagen-HA gels were significantly more swollen (Fig. 2a). While keeping the collagen concentration fixed at 2.5 mg/mL, we varied the amount of HA in the gel from 0.1 to 2.0 mg/mL. We found that the swelling ratio of collagen-HA gels increased from 1.06 ± 0.07 to 1.14 ± 0.01 as the concentration of HA increased. Using the free swelling ratio measured at different HA concentrations, and the shear stiffness (next section), we determined the model parameters *α, μ_col_, κ_col_, μ_GAG_, κ_GAG_* as described in SI sec 4. The dependence of the swelling ratio on the HA concentration also agrees well with studies on tumor swelling, which showed that tumors were no longer swollen after degradation of GAGs^12^. We note that we mixed collagen and HA (1.5 MDa) without crosslinking agents when preparing the gels. The collagen-HA co-gels should only contain physical entanglements between collagen and HA without chemical crosslinks. A fraction of HA diffused out of the collagen network during swelling (Fig. S4). Still, the HA constrained in the collagen network caused the gels to swell. The swelling observed in our experiments thus suggests that the collagen network can physically restrain movement of large HA molecules.

**Figure 2.**
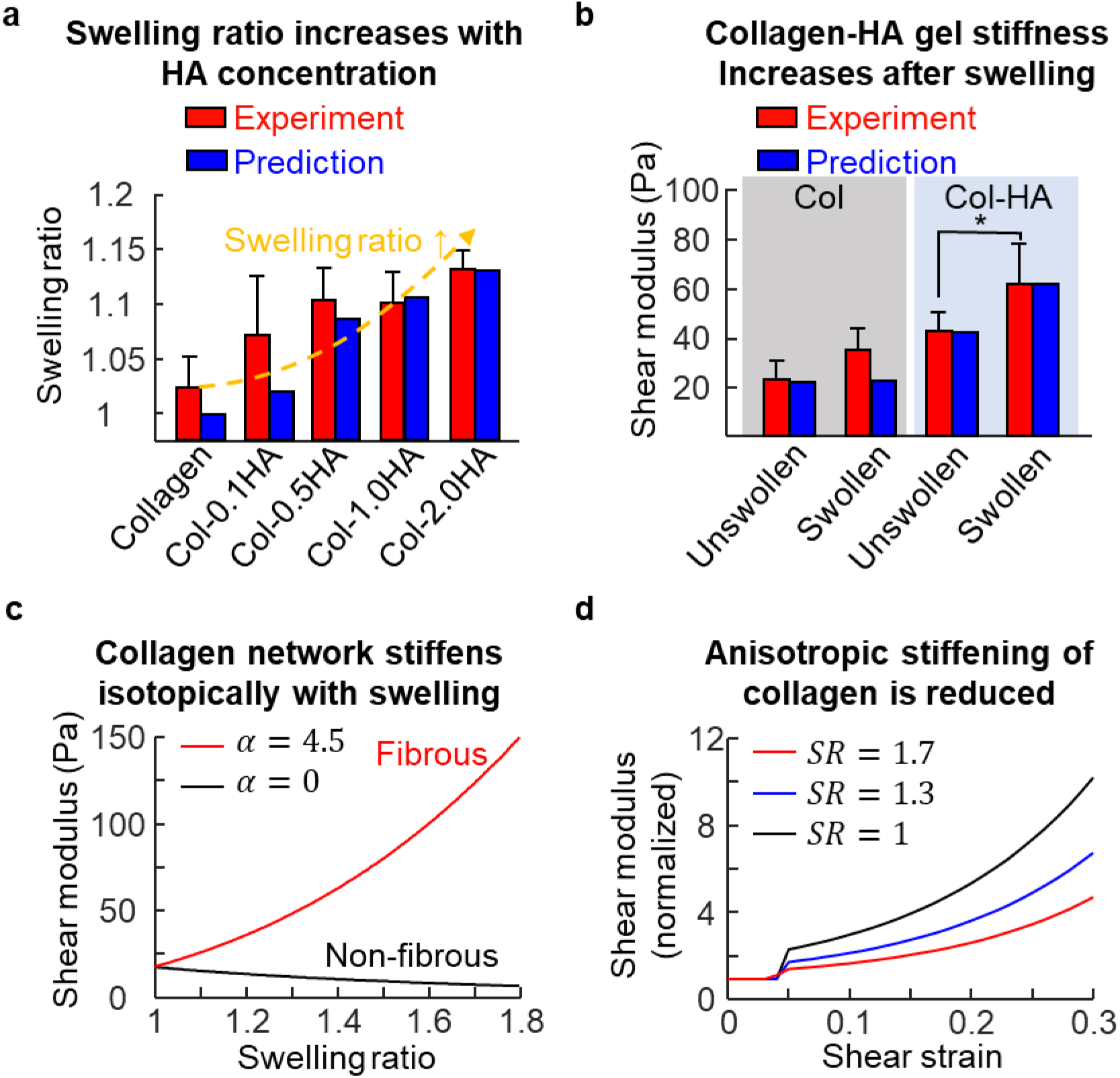
Collagen (Col; 2.5 mg/ml) gels with varying amounts of HA (1.5 MDa, 0.1-2.0 mg/ml). (a) The change in gel volume following 24 hours of submersion in pure water. An increase in HA concentration increases the swelling ratio, which is captured by the theoretical model. (b) The gel shear modulus (measured at 1% strain) before and after swelling. (c) The predicted change in matrix shear stiffness with swelling ratio. (d) The predicted change in the matrix shear modulus with shear strain, normalized to the instantaneous shear modulus.

### Collagen network stiffens isotopically due to swelling induced by GAGs

Next, we investigated how swelling can change the stiffness of the collagen-GAG network. Our discrete fiber simulations revealed that when GAGs attract water to expand the collagen network, 1) the overall spatial density of collagen decreases, and 2) the fibers strain-stiffen upon stretching of the initially wavy structures (Fig. 1b). The former causes a decrease of the network stiffness, similar to the effect of decreasing stiffness of entropy-dominated polymer networks such as polyacrylamide as they swell^23^. This mechanism competes with stiffening from the straightening of individual fibers, and the ECM can either stiffen or soften depending on which mechanism is dominant. Our continuum model captured these behaviors well: the elasticity of the GAGs (Eq. 10) and the unaligned fibers (Eq. 12) is described by the non-fibrous constitutive law (Neo-Hookean), which softens upon volume expansion. On the other hand, the scaling factor *s*(*J*), determined from discrete fiber simulations, captures the stiffening effect (Eq. 13). Our model predicted that if the impact of straightening of collagen fibers is small (i. e. *α* ≈ 0 in the model), the stiffness of the network decreases upon swelling. On the other hand, when *α* (Eq. 13) is large, the collagen-GAG network transitions from softening to stiffening behavior upon swelling (Fig. 2c).

To identify whether the collagen-GAG network stiffens in physiologically relevant scenarios, we measured the storage (shear) modulus of the collagen-HA gels (see materials and methods section for details). We fixed the collagen concentration at 2.5 mg/mL and HA concentration at 2 mg/mL, and let the gels swell freely for 24 hours at room temperature. Before swelling, the collagen-HA gel (35 ± 10 Pa) was clearly (p < 0.001) stiffer than the pure collagen gel (21 ± 10 Pa), indicating that HA was physically linked to the collagen network, contributing to the elasticity of the collagen-HA co-gel. This result agrees with data from the literature^5^, which shows that addition of HA can enhance the elasticity of the network. After swelling, we found that the collagen-HA gels became significantly (p < 0.001) stiffer with about a 2-fold increase in shear stiffness to 62 ± 18 Pa (Fig. 2b). This was because the initially relaxed fibers in the collagen network became straightened and stretched in the process of swelling^24^. Note that this stiffening behavior of the semi-flexible collagen network is not present in polymeric hydrogels (e. g. crosslinked HA), where stiffness is dominated by the entropy of the flexible polymeric chains. Swelling can decrease the density of entropic springs and thus lead to softening. To compare the behavior of collagen networks and polymeric hydrogels, we conducted a separate free swelling experiment with non-fibrous HA gels. Note that these HA were thiol-modified to form crosslinked gels, different from the linear HA in the collagen-HA co-gels. After gelation, the shear modulus was measured to be 243 ± 20 Pa. The stiffness decreased to 186 ± 11 Pa after swelling in PBS (swelling ratio 1.3), and further decreased to 138 ± 20 Pa after swelling in pure water (swelling ratio 3.3) (Fig. S3). Softening of polymeric gels (e. g., styrene butadiene rubber, polyacrylamide) upon swelling has also been reported in previous studies^23,25^. Therefore, in contrast to collagen gels, non-fibrous gels did not stiffen upon swelling, validating the predictions of our theoretical model when *α* = 0 (Fig. 2c).

We used the measurements of stiffness and free swelling ratio to quantitatively calibrate and validate our model for the elastic response of collagen-HA networks. We used a grid search to sweep the space of five parameters *α, μ_col_, κ_col_, μ_GAG_, κ_GAG_*, and chose the set that best fit (with lowest mean squared error) the free swelling ratio (Fig. 2a) and shear moduli before and after swelling (Fig. 2b). Other parameters were obtained from the literature (see SI sec. 4 table of parameters). With the parameters given in SI sec. 4, our model quantitatively captured 1) the increase in swelling ratio with increasing GAG concentration, and 2) the increase in network stiffness upon swelling (Fig. 2a & 2b). The excellent agreement between our simulations and experiments shows that for modest levels of swelling (*SR* = 1~1.2), the stiffening induced by straightening fibers outpaces the softening due to decrease of density, leading to an increase in matrix stiffness.

### Anisotropic strain-stiffening of collagen networks is less prominent after swelling

Next, we studied the strain-stiffening behavior of the network at different levels of swelling. Using the model parameters obtained from the free swelling experiments, we further simulated shear deformation on the network at three levels of swelling (*SR* = 1, *SR* = 1.3, *SR* = 1.7). After normalizing the shear modulus of the network with respect to the instantaneous shear modulus at zero shear strain, we compared the change of the shear modulus as a function of the shear strain. We found that as the initial swelling increases, the increase of the shear modulus is less apparent (Fig. 2d). At 0.3 shear strain, the shear modulus increases by 10 fold compared with the instantaneous shear modulus when *SR* = 1. This level of stiffening is about twice as much as the case for *SR* = 1.7, which indicates that the strain-stiffening of the network is less prominent when the network is swollen. This is because 1) the swelling increases the stiffness of the network isotopically by stretching the fibers, and 2) anisotropic stiffening of the network due to fiber alignment is less affected by the isotropic expansion of the network. With increase of the isotropic stiffness, the network becomes less non-linear, and the anisotropic stiffening is less prominent. Note that the kink in the curves (Fig. 2d) is due to the increase of network shear stiffness as fibers start to align along the directions of tension. On the other hand, we have previously identified that the strain-stiffening of the network along the direction of tension is essential to the long-range force transmission^2^. Therefore, we predict that the range of force transmission decreases with network swelling, which can be induced by GAGs. In the next section, we present the analysis of the range of force transmission as a function of GAG content in the collagen network.

### Range of cell force transmission decreases with GAG content

Having identified the role of GAGs on the mechanics of collagen co-gels, we proceeded to study how GAGs can alter the transmission of the active forces generated by cells. We first considered a simple case in which a contractile cell was embedded in a matrix with homogeneous distribution of GAGs. We modeled the cell as an ellipsoid described by the equation (*x/a*)^2^ + (*y/a*)^2^ + (*z/b*)^2^ ≤ 1, where *a* and *b* denote the half-length of the cell short and long axes (Fig. 3a & 4a). We define the cell aspect ratio, *AR* = *b/a*, which increases as the cell becomes elongated. To model the impact of cell shape (at fixed volume) on force transmission, we set 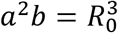. Here *R*_0_ is the radius of the cell when it assumes a spherical shape. In our previous study^2^, we determined that the isotropic mechanical behavior of the unaligned collagen fibers cannot explain long-range force transmission; forces were transmitted at distances greater than the size of the cells due to the alignment of fibers in response to cellular forces. In our simulation setup, we let the ECM first swell freely under physiological conditions with 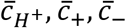 on the outer boundary at 0.0001 mmol/L (pH = 7), 150 mmol/L, 150 mmol/L, respectively. We then imposed a prescribed displacement boundary condition at the cell-ECM interface to simulate the contraction of the cell. Specifically, the cell contractile strains along the *x, y, z* directions (*ϵ_x_, ε_y_* and *ϵ_z_*, respectively) were set to be 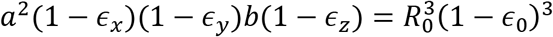, so that the cell volume after contraction was the same for all shapes. Here *ϵ*_0_ is a constant and *ϵ_x_* = *ϵ_y_* = *ϵ_z_* = *ϵ*_0_ when the cell is spherical (*AR* = 1). As shown by previous studies^3^, the contraction of the cells becomes larger along their long-axis as they become elongated, and therefore we assumed *ϵ_z_*/*ϵ_x_* = *ϵ_z_/ϵ_y_* = *AR*, so that the contractile strain was larger along the long axis for elongated cells. The outer boundary of the ECM was prescribed such that no displacement was allowed in directions perpendicular to the boundary. We ensured that the overall size of the simulation box was sufficiently large to eliminate boundary effects (matrix boundary to cell distance ≫ 40*R*_0_). We repeated the simulations for various concentrations of GAGs and studied the range of force transmission.

**Figure 3.**
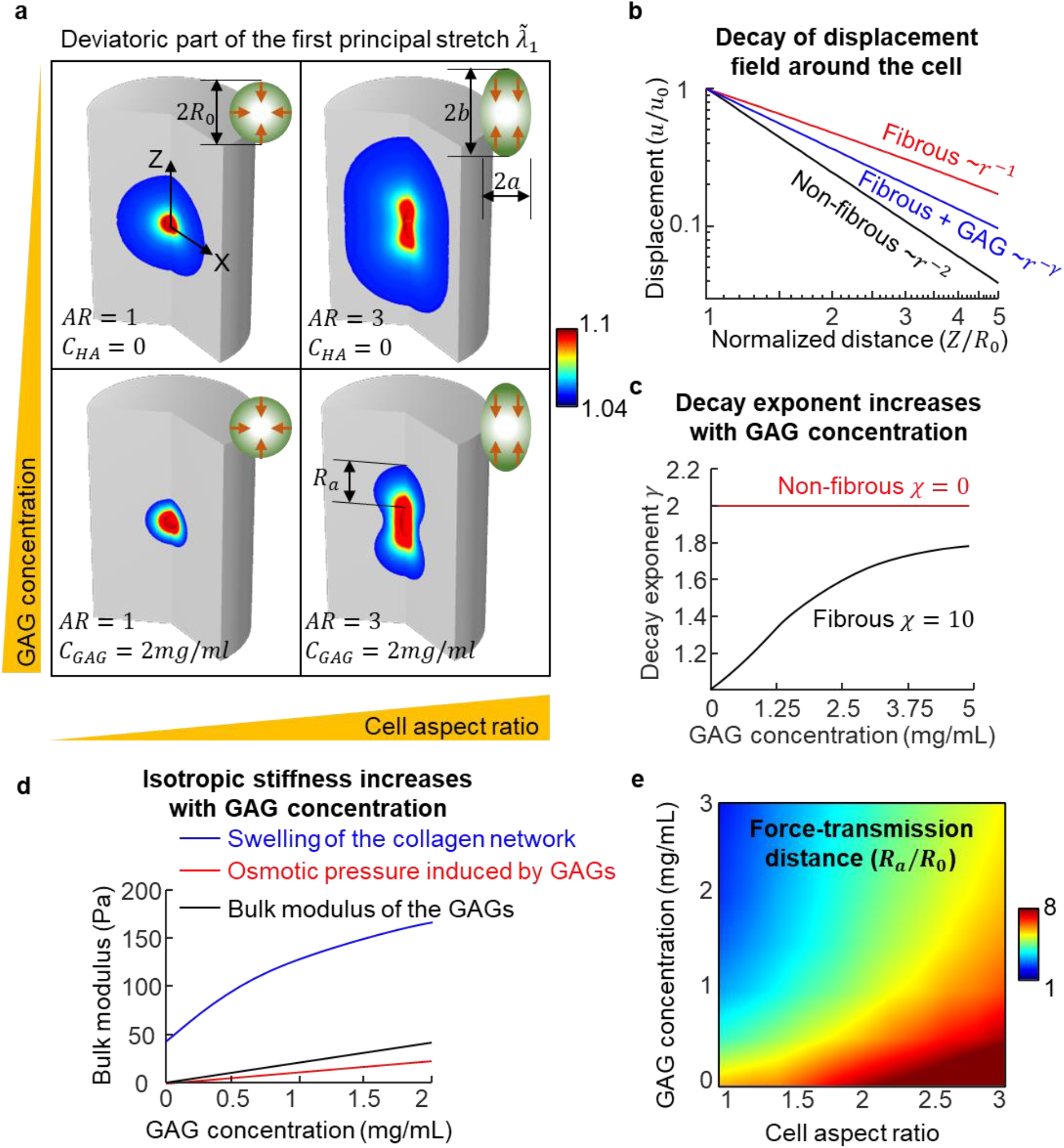
(a) Influence of cell shape and GAG concentration on the distance (*R_a_*) over which the cell-induced contractile force is transmitted measured by the extent of the regions where fibers are aligned in matrices. We define aligned regions as part of the matrix with the deviatoric part of the first principal stretch larger than the critical value 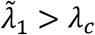. (b) The decay of matrix displacement as a function of the distance from the cell. (c) Dependence of the decay exponent on GAG concentration. (d) Contribution to isotropic stiffness of the collagen-GAG co-gels from 1) swelling of the collagen network, 2) the bulk modulus of the GAGs and the osmotic pressure induced by the GAGs. (e) Heat map of the range of force transmission, *R_a_*, as a function of cell shape and GAG concentration.

**Figure 4.**
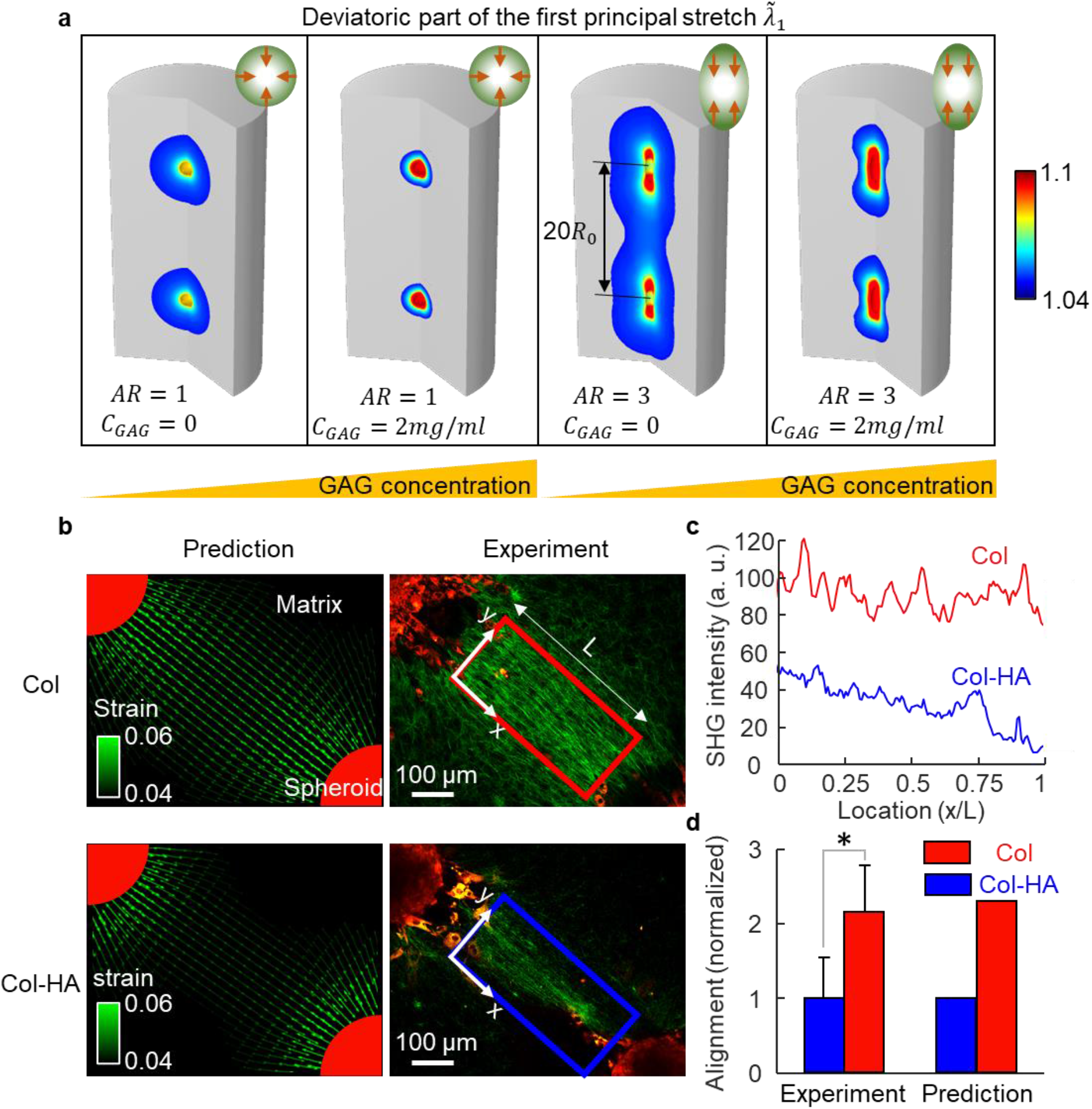
(a) Theoretical prediction of fiber alignment between two cells in collagen-GAG co-gels with increasing GAG concentration. (b) Theoretical prediction and experimental measurement of alignment of collagen fibers between cell spheroids. (c) The SHG intensity averaged along the y-direction plotted as a function of the location along the x-direction specified in (b). (d) The average fiber alignment in the box specified in (b), n=17 for collagen gels and n=9 for collagen-HA gels, p < 0.001.

With numerical simulations, we found that for a given level of cell contraction, there was a significantly decreased level of strain in the vicinity of the cell when GAGs were present (Fig. 3a) regardless of the shape of cell. In the case when the cell is spherical, the decay of cell-induced displacement field *u*(*r*) was found to follow a power law *u*(*r*)~*r*^−*γ*^, in which *r* denotes the distance to the center of the cell^2^. In this case, we were able to derive an analytical expression for the decay exponent,

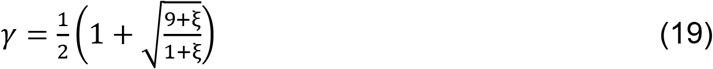

where 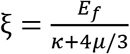 in which 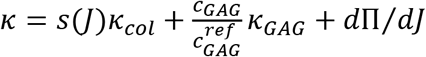 and 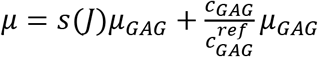 are the isotropic bulk and shear modulus of the GAG-collagen network that include contributions from both the GAGs and unaligned collagen fibers. When there are no GAGs in the network, 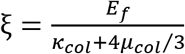, where *E_f_* (characterizing the anisotropic stiffening of the network) is much larger than the isotropic stiffness *κ_col_* and *μ_col_*. In this case, *ξ* approaches infinity and the exponent *γ* approaches 1, indicating a slow decay of displacement. On the other hand, *γ* = 2 when the network is not fibrous and thus does not strain-stiffen in an anisotropic manner (*E_f_* = 0, *ξ* = 0). When GAGs are present in the network, the isotropic stiffness *κ* and *μ* increase because GAGs directly contribute to the isotropic stiffness (captured by *κ_GAG_* and *μ_GAG_*) and induce swelling that stretches and stiffens the collagen network (captured by the scaling factor *s*(*J*)). Additionally, compressing (stretching) the network leads to a higher (lower) concentration of fixed charges, and thus a difference in osmotic pressure captured by *d*Π/*dJ*, which contributes to the bulk stiffness of the network^26^. The contributions of these three mechanisms to the isotropic stiffness as a function of the GAG concentration is shown in Fig. 3d. We found that the contribution from the stiffening of the collagen network due to the swelling induced by GAGs was significantly larger than the other two mechanisms. With higher isotropic stiffness *κ* and *μ* in the presence of GAGs, the decay exponent *γ* falls in between 1 and 2, with an increase in the value of the decay exponent with the concentration of GAGs as shown in Fig. 3c.

Since the cell-induced displacement decays faster when GAGs are present in the network, we also predict a decrease in the range of force transmission. Through our numerical simulations, we found the furthest distance that the cell-induced force can reach (*R_a_*, defined by 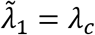) decreased with increasing GAG content in the network (Fig. 3a). To quantitatively investigate *R_a_*, we conducted an analytical analysis (SI sec. 2), and found that for a spherical cell, *R_a_* can be calculated by

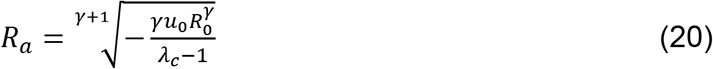

Here, *R*_0_ denotes the radius of the cell, and *u*_0_ = *ϵ*_0_*R*_0_ denotes the radial displacement at the cell-matrix interface. From this equation, it is apparent that the range of cell force transmission increases when the cell is more contractile (which can exert a higher displacement *u*_0_). For the same level of contraction *u*_0_, *R_a_* decreases with the decay exponent *γ* (shown in SI sec. 2). Therefore, we predict that the range of force transmission decreases with the concentration of GAGs as shown in Fig. 3a. Overall, high GAG concentration increases the isotropic stiffness of the collagen network, and makes the anisotropic strain-stiffening less prominent. The network behavior approaches that of a non-fibrous material (such as a linear elastic or Neo-Hookean material), in which the cellular force decays rapidly.

We then examined how the range of force transmission depends on GAGs when cells are elongated. Due to the anisotropy of the cell geometry and contractility^2^, the contraction of an elongated cell caused a “gourd”-shaped region of fiber alignment (Fig. 3a). The elongated cell with an aspect ratio of 3 can transmit force over three times further than a round cell in a collagen matrix (Fig. 3a). Similar to the case of a round cell, when we increased the GAG concentration from 0 to 2 mg/mL, we found a significantly reduced range of force transmission. In Fig.3e, we show the range of force transmission as a function of both GAG concentration and cell aspect ratio. For all the cell shapes that we investigated, GAGs reduce the range of cellular force transmission. A more comprehensive parametric study on the relation between the range of force transmission and other parameters (*χ, λ_c_*, and *u*_0_) is included in SI sec. 2. We find that the range of force transmission increases when 1) the collagen network strain-stiffens more, 2) the collagen network has lower critical strain for alignment, and 3) the cell exerts larger levels of contractile forces (Fig. S1).

### GAGs suppress long-range cell-cell communication

Interactions between cells are involved in many biological processes (e.g., cancer cell invasion, wound healing, and fibrosis) and play a significant role in the organization of tissues. Since the range of force transmission decreases in the collagen network with GAGs (as presented above), it is reasonable to expect that cell-cell interactions will also be impacted. To verify this hypothesis, we explicitly modeled the interaction between two contracting cells embedded in collagen matrices with varying amounts of GAGs. As in the case of single cells, where the deformation fields spread further from the cells when they are elongated, elongated cells interact more significantly compared to spherical cells (Fig. 4). In the matrix without GAGs, we observed a significant overlap of regions of alignment between the two cells at a distance of 20 times *R*_0_ (Fig. 4a). The aligned regions were stiffened anisotropically along the line connecting two cells, allowing efficient transmission of forces between the cells. However, with the addition of GAGs to the matrix, the regions of alignment no longer overlapped, indicating that the contraction of one cell cannot be felt by the other cell (Fig. 4a). Using these simulations, we confirmed that the cell-cell interaction distance was twice the size of the cell force transmission range, indicating that increasing GAG content can decrease long-range cell-cell communication.

To validate the prediction on the range of cellular force transmission, we studied the alignment of collagen fibers between two fibroblast spheroids. Since the cell aggregates remained roughly spherical (Fig. 4b) during the experiment, our model that considered two single spherical cells can be used here by treating each spheroid as a large single source of contractile force, which can be analyzed using the numerical and analytical methods shown in model development section and SI sec.2. Fibroblast spheroids were seeded 600 μm apart on a collagen type I gel (2.5 mg/mL), or collagen-HA co-gel (2.5 mg/mL collagen and 2 mg/mL HA. The gels were formed on coverslips and exposed to the culture medium, after which fibroblasts began pulling the matrix following the formation of adhesions. Within a few hours, the contraction led to large-scale fiber alignment, especially in the region connecting the two spheroids (Fig. 4b). We used second-harmonic generation (SHG) microscopy to visualize and image aligned collagen fibers. We extracted the average SHG intensity along the y-direction and plotted it along the x-direction (Fig. 4b & 4c). We found that the collagen alignment was much less prominent between cells cultured on the collagen-HA gel, despite the fact that fibroblasts should be more contractile on the stiffer collagen-HA substrate^27^. This result suggests that GAGs decrease long-range force transmission in the matrix and agrees very well with the alignment of collagen fibers predicted by our model. We adopted the parameters determined from free swelling and rheology experiments (parameter values are listed in SI sec. 4), and simulated the contraction of fibroblasts with 10% contractile strain. We visualized the deviatoric part of the first principal strain 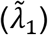, whose magnitude indicates the density of aligned collagen, as shown in Fig. 4c. We found a quantitative agreement between the SHG intensity and the magnitude of the principal strain for both collagen and collagen-HA matrices (Fig. 4d). The excellent quantitative agreement between model predictions and experiments confirms that the presence of GAGs indeed significantly modulates the range of force transmission in collagen matrices.

## Discussion

In this study, we developed and validated a theoretical model describing the mechanical behavior of collagen networks with interpenetrating GAGs. By combining predictions and experiments, we showed that GAGs can attract water into the ECM, stretching the collagen fibers and leading to an increase of network isotropic stiffness, and that the ECM becomes more resistant to volumetric changes. Collagen fibers stretched under swelling can better resist buckling and were less likely to align along the direction where the stresses were tensile (Fig. 5). For the same level of shear load, the anisotropy of the fiber orientation decreased with the swelling ratio. By preventing the reorientation of collagen fibers that is essential to long-range force transmission, GAGs decreased long-range force transmission. In agreement with our model predictions, we found significantly decreased collagen alignment between cell spheroids in collagen-HA co-gels compared with collagen gels. We further showed that the local accumulation of GAGs can create barriers to cellular force transmission (Fig. S2). This is physiologically relevant because the distribution of GAGs can be highly inhomogeneous *in vivo*. For example, in breast tumors, high concentrations of GAGs are only observed at the leading edge of the tumor^7^, and the localization of GAGs has been linked to the severity of cancer.

**Figure 5.**
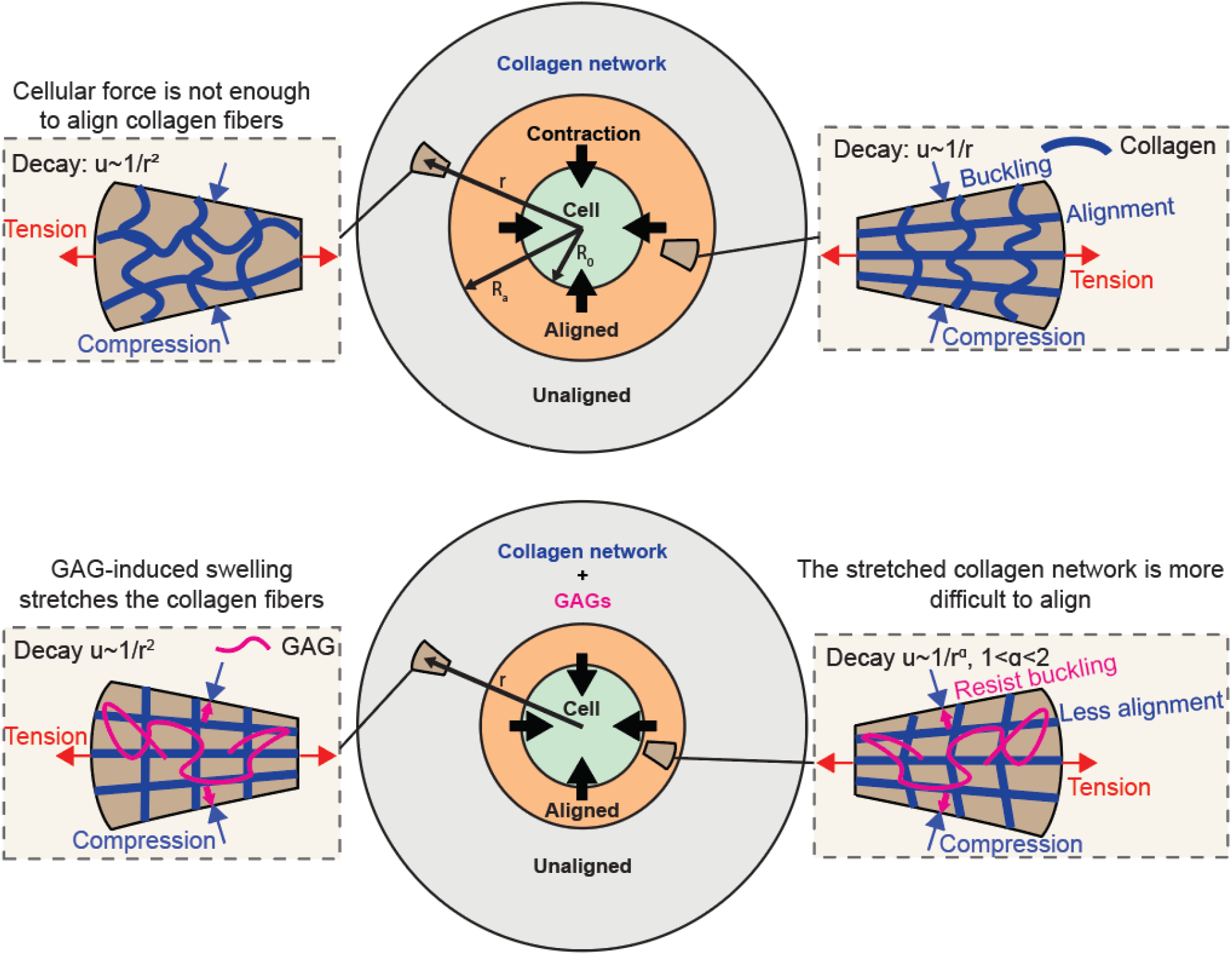
Schematic showing the role of GAGs in cellular force transmission. In the matrix without GAGs (top), contraction of the cell aligns the fibers in the radial direction. In the presence of GAGs, collagen networks are better able to resist compression and buckling in the transverse direction. This mechanism reduces collagen fiber alignment in the radial direction and reduces the efficiency of force transmission.

While we experimentally validated our model using HA as the model GAG, the ECM *in vivo* also contains other types of GAGs such as chondroitin sulfate and heparan sulfate. Often, GAGs attach to a core proteins to form proteoglycans, such as the highly GAG-modified bottle-brush aggrecan, which is further bound to HA chains to form a larger proteoglycan complex^28^. Like HA, these GAGs contain acidic groups (e.g., carboxyl, sulfate) that can dissociate into fixed negative charges and induce swelling with the same mechanism^9^. Therefore, the theoretical predictions in this study can be applied to other GAGs that can attract water and swell the matrix. However, in native ECM, the content of free GAGs and PG-bound GAGs is still unclear. As different PGs appear to interact differently with collagen^29^, PG-bound GAGs (fixed charge residues) might have complicated effects on mechanical communication between cells. Thus, there is a particular need to investigate and compare the regulation of free versus PG-bound GAGs on fibrous network as one future direction for modeling force transmission in native ECM.

Our results have implications for cellular mechano-transduction in biological systems where GAGs are involved. For example, cancer-associated fibroblasts (CAFs) behave very differently than their normal counterparts in ways that allow them to thrive and contribute to the development of the tumor microenvironment^9^. One of the important distinctions between the two kinds of fibroblasts is that CAFs have a significantly increased ability to synthesize HA^30^. GAGs are also abundant in connective tissues such as cartilage and tendon. These tissues undergo constant mechanical loading, and GAGs are critical to maintain the hydration necessary for them to function normally. Damaged cartilage usually shows decreased GAG content. Our model provides a foundation for the analysis of cell mechano-sensing in these scenarios, and can potentially help develop treatments for disease conditions related to changes in GAG content^31^.

In this study, our focus has been on the impact of GAGs on cellular force transmission. However, GAGs also play a role in mechanosensing by attaching to cell membrane receptors such as the HA receptors CD44 and RHAMM (Fig. S2a). GAGs are a major component of the pericellular matrix (PCM), also known as the glycocalyx^8^, which can extend as much as 20*μm* from the cell surface. An osmotic pressure gradient has been experimentally observed^32^ in the PCM as the density of GAG chains increases approaching the cell surface, and may be relevant in cell mechanosensing. For example, several studies have shown that HA in the PCM can have both adhesive^33,34^ and antiadhesive^35–37^ roles. Surface HA is also involved in transducing shear stress to endothelial cells^38^. HA upregulates the contractility of periodontal ligament cells via its interaction with CD44^39^. We hope to further investigate these roles of GAGs in forthcoming work.

## Materials and Methods

### Gel preparation and free swelling experiments

Collagen-HA gels were prepared by mixing rat tail collagen I (Corning) to a final concentration of 2.5 mg/mL with a varying amount of HA (collagen to HA weight ratio: 2.5:0.1,2.5:0.5, 2.5:1, and 2.5:2, 1.5 MDa sodium hyaluronate from Lifecore Biomedical). The gel solutions were incubated at 37°C overnight and were then submerged in deionized water at room temperature, subject to no force. After 24 hours, we wiped excessive water from the gel surface and measured the weight. The swelling ratio (*SR*) was then calculated by

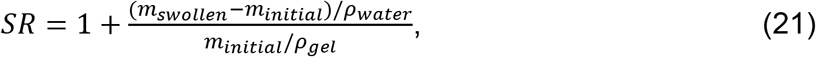

where *m_initia[_* and *m_swollen_* denote the weight of the gel before and after swelling, respectively. *ρ_water_* and *ρ_gel_* denote the density of water and gel prior to swelling, respectively. As the mass concentrations of collagen and hyaluronan are less than 1%, we assume that *ρ_gel_* ≈ *ρ_water_*. Note that the HA used here is not crosslinked and only physically restrained in the collagen network. After 24 hours, almost 50% of hyaluronan has diffused out of the gel (Fig. S5). However, the HA that remains in the collagen gel network still induced swelling compared to the pure collagen (Fig. 2a). Crosslinked HA hydrogel was made by mixing Glycosil (thiol-modified HA from Advanced BioMatrix) with Extralink (thiol-reactive crosslinker, polyethylene glycol diacrylate (PEGDA)) at a volume ratio of 4:1 and formed on a 20mm coverslip. The swelling experiments were carried out as described above.

### Shear rheology experiments

For rheometry of collagen, collagen-HA and crosslinked HA gels, the gel solutions were prepared as previously described for the free swelling experiments. Unswollen gels were measured after gelation and swollen gels were measured after swelling in deionized water for 24h. A shear rheometer (Kinexus) with rSpace software was used to characterize shear modulus. A 20mm plate was used and both storage and loss (G’ and G”) were measured by applying an oscillatory shear strain of 2% at a frequency of 10 rad/sec (1.592Hz).

### Fibroblast spheroid experiment

Collagen and collagen-HA gels were prepared as described for free swelling experiments using microwell plates with glass-bottomed cutouts. Plates were sealed with parafilm and kept in an incubator overnight at 37°C. Fibroblast spheroids were formed by the hanging droplet method. Briefly, passaged rat portal fibroblasts^40^ were trypsinized and suspended in culture media at 200,000 cells/ml. 20 μL droplets of suspension cell solution were placed on the underside of a dish lid. To avoid drying, 10 ml media was added to the dish. After inversion of the lid, the cell droplets were cultured for 3 days. For seeding, spheroids were captured and carefully placed on the gel in pairs approximately 600 μm apart. The gel was incubated for 4h allowing the spheroids to attach and then 2 ml media was added. After culturing for another 20 h, gels were rinsed with PBS and fixed with 10% formalin for 10 min. All the samples were sealed with parafilm and kept in PBS at 4°C. Second-harmonic generation (SHG) imaging was used to visualize collagen fibers^29^.

### Statistical analysis

The significance of changes in shear moduli and swelling ratio were tested using one-way ANOVA; SHG intensities were compared using student’s t-test (unpaired). p value was chosen to be 0.001.

### Discrete three-dimensional fiber network model

Three-dimensional fiber networks were generated by placing fibers over the edges of a three-dimensional Voronoi tessellation. The Voronoi diagrams were generated by random seed points using MATLAB (MathWorks, Natick, MA). The individual fibers were modeled as Timoshenko beams^41^, flexible in bending, stretching, twisting, and shear modes of deformation. Fiber waviness was introduced by shaping each fiber into a half-sine wave. The amplitude of the sine waves equaled half of the fiber end-to-end length. The fibers had circular cross-sections of diameter 150 nm and elastic moduli of 6.5 MPa. Implicit finite element calculations were performed using the Abaqus software package (Simulia). The shear tests were performed by the horizontal displacement of the top surface, in the x direction, while the bottom was held fixed. The shear tests of the expanded networks were performed in two steps. In the first step, the volumetric expansion of the networks was modeled by the stretching of the networks equally in three directions by enforcing displacements at the boundaries. The reaction forces at the boundary nodes over all surfaces except the top and bottom surfaces were recorded at the expanded state using the Abaqus scripting interface in Python. In the second step, displacement boundary conditions were applied to the nodes on the top and bottom surfaces, while the recorded reaction forces due to volumetric expansion were applied to the nodes on all faces except for the top and bottom faces. Fiber reorientation was characterized by the measurement of the distribution of the orientation of fibers stretched by more than 1%. The orientation of individual fibers was evaluated by the projection of the fiber’s end-to-end vector onto the front surface of the network and the calculation of the orientation of the projected vector with respect to the x-direction.

## Acknowledgments

This work was supported by National Cancer Institute awards R01CA232256 (VBS), U54 CA261694 (VBS), National Institute of Biomedical Imaging and Bioengineering awards R01EB017753 (VBS, RGW, PJ) and R01EB030876 (VBS), NSF Center for Engineering Mechanobiology grant CMMI-154857 (RGW, PJ, VBS), NSF grants MRSEC/DMR-1720530 (RGW, PJ, VBS) and DMS-1953572 (VBS).

## Author Contributions

X. C., E. B., and V. B. S designed the theoretical models and carried out the computations; D. C., R. G. W, and P. A. J designed and conducted the experiments; X. C., D. C., E. B., R. G. W, P. A. J, and V. B. S analyzed and interpreted the data; X. C., D. C., E. B., P. A. J, R. G. W, and V. B. S wrote the paper.

## Supplementary information

### 1. Analytical analysis of the impact of GAGs on the cellular force transmission

To derive an analytical expression for the range of force transmission, we considered a spherical cell (of radius *R*_0_) exerting isotropic contractile stress on the surrounding swollen matrix (Fig. 5). Choosing the swollen state as the reference, the strain is tensile (compressive) in the radial (transverse) direction. In the vicinity of the cell, the radial strain induced by cell contraction is large enough to start to align fibers in the matrix. Due to spherical symmetry, the region where the fibers are aligned is also spherical with radius *R_a_*. Inside this region, we can write the matrix stress as

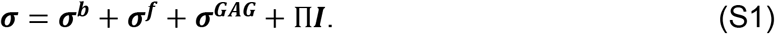

Here, ***σ^b^, σ^f^, σ^GAG^**, and Π***I**** denote the stress tensors that arise from the randomly-aligned collagen, aligned collagen, GAGs, and the swelling pressure induced by GAGs. Inside the aligned region, for linear bulk and fibrous response (*m* = 0 in Eq. 15),

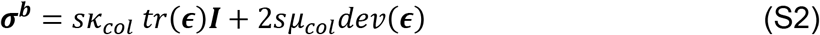

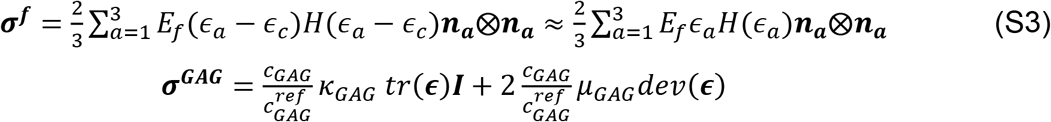

Here, *H*() denotes the Heaviside step function, and *ϵ_c_* = *λ_c_* – 1 ≪ 1 denotes the critical strain for the collagen matrix to align. Note that in the swollen configuration the bulk modulus *κ* and shear modulus *μ* are scaled by the scaling factor *s*(*J*) as shown in Eq. S2. Using a spherical coordinate system (*r, θ, φ*), we write the strains as

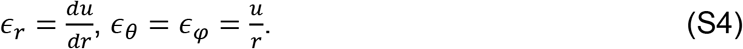

Eq. S1 can then be rewritten as

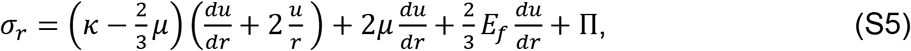

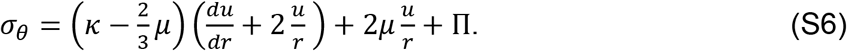

Here in these equations, 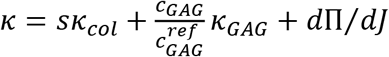 and 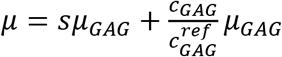 denote the effective isotropic bulk and shear moduli of the network. The condition for mechanical equilibrium 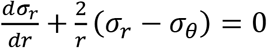 can then be written as,

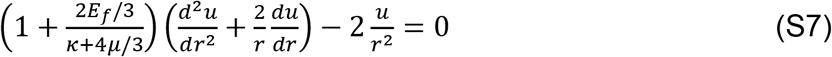

with boundary conditions

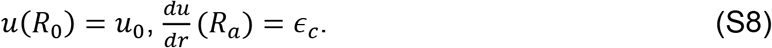

Here *u*_0_ denotes the displacement at *r* = *r*_0_ induced by cell contraction. When *ϵ_c_* ≪ 1, the solution can be approximated by

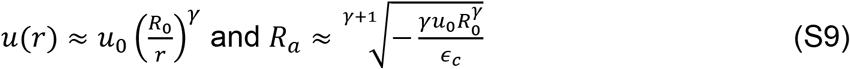

where 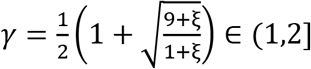 and 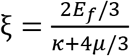. When the matrix has no fibrous component with *E_f_* = 0, *ξ* = 0, *γ* = 2, the displacement decays fast as 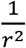. On the other hand, when the matrix is highly fibrous with *E_f_* → ∞, *ξ* → ∞, *γ* → 1, the displacement decays much more slowly as 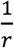, showing a long range of force transmission.

For the region *r* > *R_a_*, the collagen fibers are not aligned and therefore the matrix stress ***σ*** has contributions only from GAGs and the unaligned fibers, i.e.

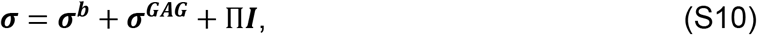

The expression of displacement in this region *u*(*r*) can be derived following a similar approach as demonstrated in Eq. S1–S9, with the boundary condition 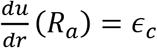 and *u*(∞) = 0. The solution follows the same form as shown in Eq. S9 with *ξ* = 0 and *n* = 2. Therefore, the overall displacement field can be written as

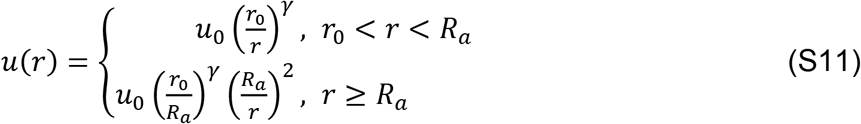

Combining Eq. S2, S3, and S11, we can derive the stress field. Here we write out the expression for *σ_r_*,

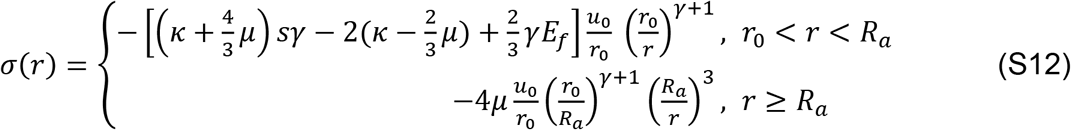

From Eq. S12, we can see that the force transmission is efficient in the aligned region *r*_0_ < *r* < *R_a_*, with stress decaying as 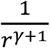. Outside the aligned region, stress decays much faster with 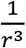.

Rescaling *u*_0_ and *R_a_* by the radius of the cell *r*_0_, we can write the normalized radius of the alignment region as 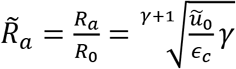, where 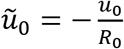. From this expression we can see 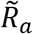 depends on *γ*, which characterizes the rate of decay of displacement in the matrix. Small (large) *γ* corresponds with slow (fast) decay of displacement, and thus long (short) range of force transmission. Since 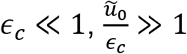, and

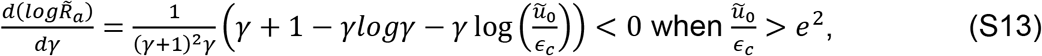

the radius of the aligned region also increases as *γ* decreases.

### 2. Relation between range of force transmission and collagen network properties

In the main text we presented the impact of GAG concentration on force transmission. Here, we show a comprehensive parametric study on the properties of the collagen network, as well as the magnitude of the cell contraction. As shown in Fig. S1, the range of force transmission increases with the magnitude of cell contraction *ϵ*_0_, and the matrix fibrous modulus 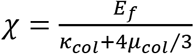.

**Figure S1.**
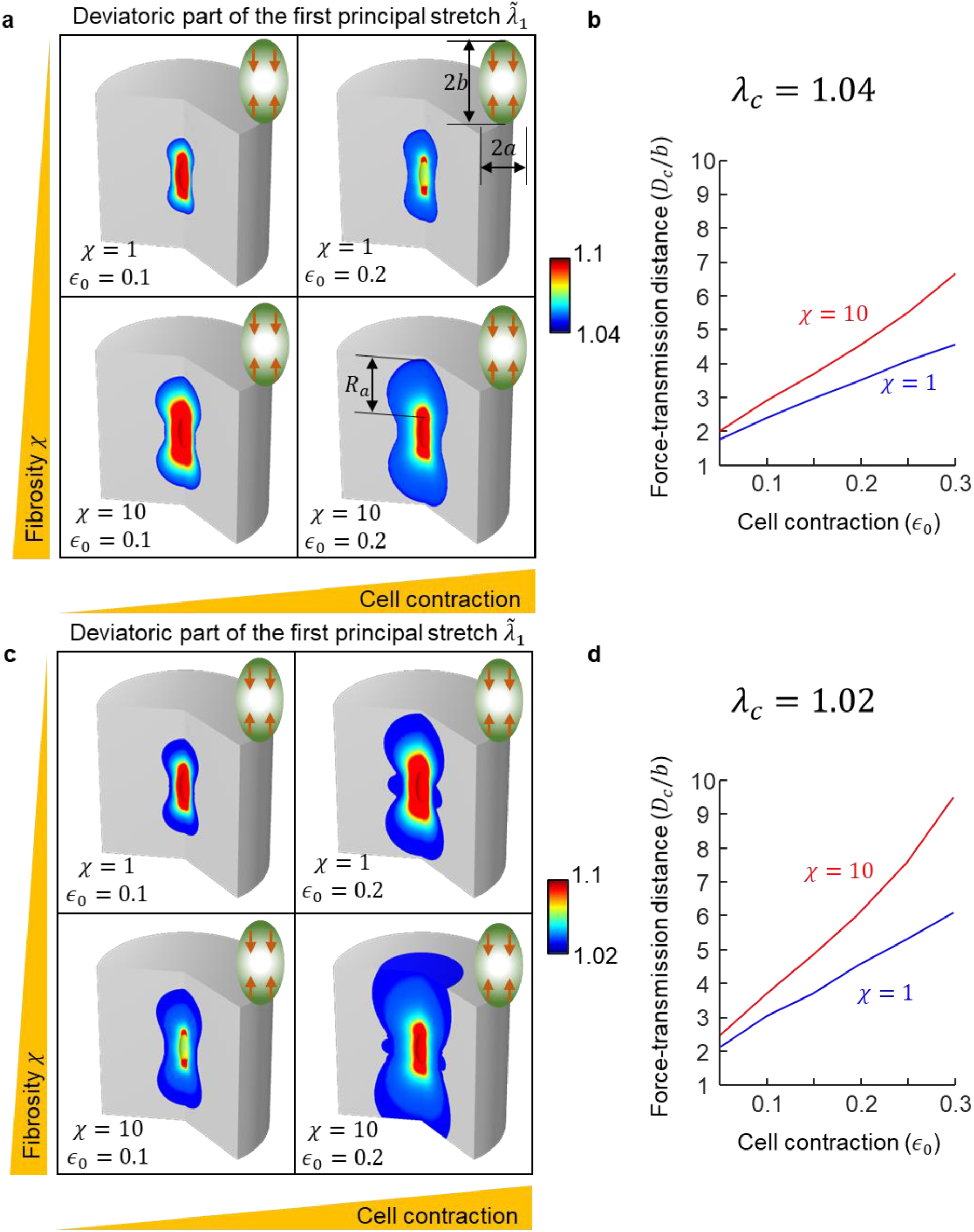
(a) Impact of cell contraction *ϵ*_0_ and matrix fibrosity *χ* on force transmission. The colored area denotes the region of fiber alignment 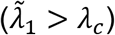. (b) Force-transmission distance normalized by the long-axis radius of the cell as a function of cell contraction *ϵ*_0_. (c) (d) Same as (a) and (d) but with *λ_c_* = 1.02.

### 3. Local accumulation of GAGs creates barriers for cell mechanosensing

Distribution of GAGs is often inhomogeneous *in vivo*. For example, in the tumor microenvironment, stromal cells have been found to release abnormally high amounts of HA, which can potentially affect cellular force transmission around the cancer cells. To investigate the impact of such local accumulation of GAGs, we simulated the increase of the concentration of GAGs in a circular region of radius *R_a_* located at a distance *L_a_* from a cell (or spheroid), defined by *x*^2^ + *y*^2^ + (*z* – *L_a_*)^2^ ≤ *R_a_* (Fig. S2b). We show that the accumulation of GAGs can disrupt the force transmission locally.

When there is no overproduction of GAGs, cell contraction aligned matrix fibers in a gourd-shaped region (Fig. S2b), consistent with our results in the main text. As GAGs start to accumulate in the small circular region (increasing *c_GAG_*), we found that within the region of GAG accumulation, the strains significantly increase because of the swelling (Fig. S2b–S2d). Normally, the cell-induced strain is tensile in the radial direction, and compressive in the transverse direction. With the presence of GAGs, the strain state is overridden by swelling. This suggests that cells located in the accumulated region cannot receive the mechanical signals from the other cells in the surrounding, which can potentially alter their mechanosensitive behaviors.

**Figure S2.**
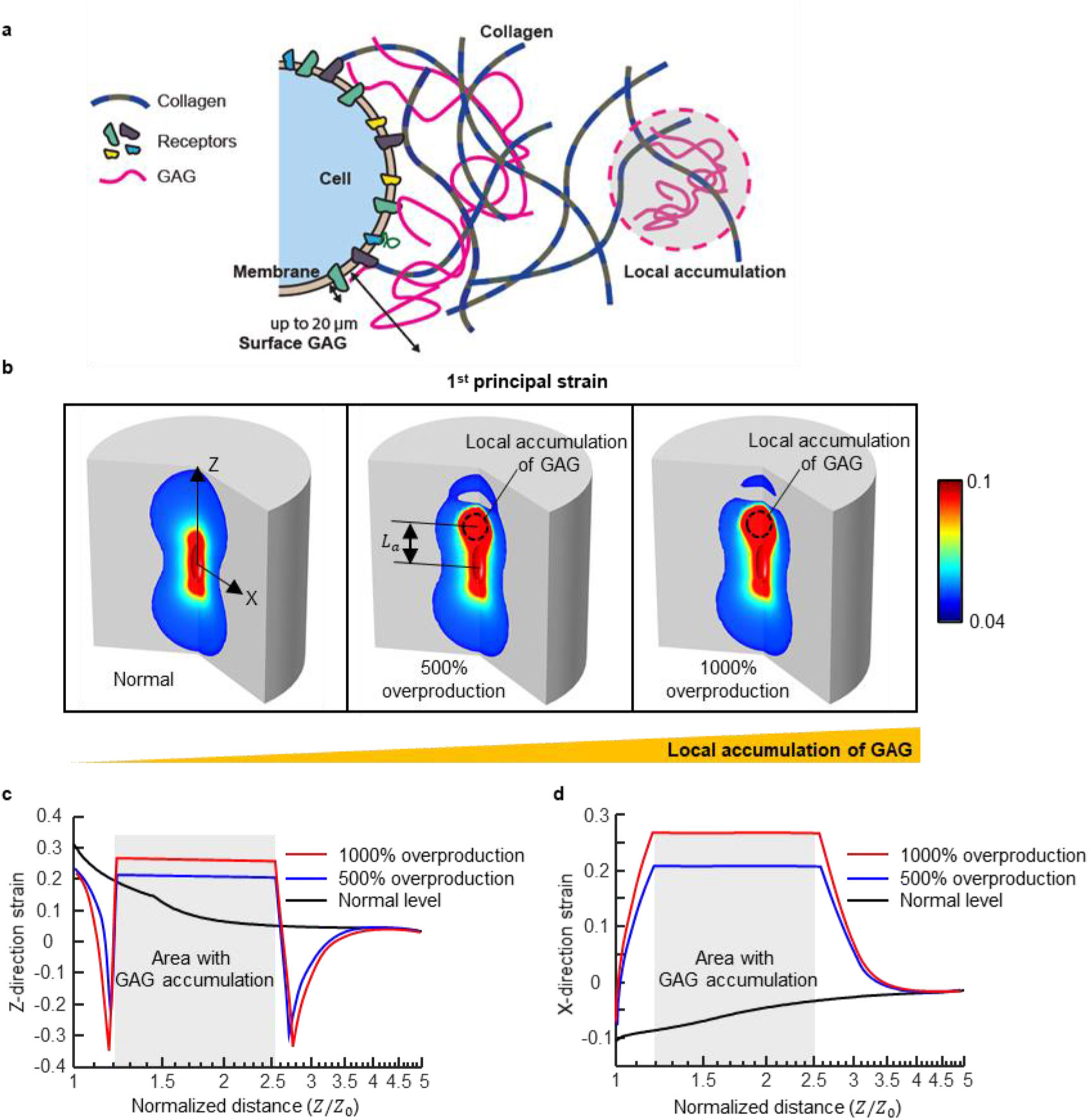
(a) Schematic showing the impact of local accumulation of GAG on force transmission. As the over-synthesized GAG leaves the cell surface and moves into a region close to the cell, it increases the local osmotic pressure and causes the collagen network to swell. The swollen region compresses the surrounding collagen network and disrupts the normal force transmission. (b) The deviatoric part of the first principal stretch in the ECM under 20% contraction of the cell. Only the region with strain larger than 0.04 is shown, indicating matrix that becomes aligned under the cell contraction. As the local accumulation of HA increases, it creates a barrier for transmission of the cell contractile force. (c) The Z-direction strain and (d) X-direction strain along the cell long axis passing through the area with increased level of GAGs.

### 4. Table of parameters

*μ_GAG_, κ_GAG_, μ_col_, κ_col_* and *E_f_* are determined by fitting the model to match the free swelling ratio and the shear moduli measured by rheology.

| Parameter | Physical meaning | Value | Source |
| --- | --- | --- | --- |
| $c_{GAG}$ | Concentration of GAGs | 0~2.5 mg/mL | 6, 7 |
| $c_{GAG}^{ref}$ | Reference concentration of GAGs for measuring stiffness | 1 mg/mL | Chosen |
| $\mu_{GAG}$ | Shear modulus of GAGs at $c_{GAG}^{ref}$ | 60 Pa | Fit to experiment |
| $\kappa_{GAG}$ | Bulk modulus of GAGs at $c_{GAG}^{ref}$ | 50 Pa | Fit to experiment |
| $\bar{c}_{H^+}$ | Concentration of hydrogen ions in the external solution | 0.001 mmol/L | Physiological condition |
| $\bar{c}_+$ | Concentration of cations in the external solution | 150 mmol/L | Physiological condition |
| $\bar{c}_-$ | Concentration of anions in the external solution | 150 mmol/L | Physiological condition |
| $K_a$ | Acidic strength | $10^{-3.3}$ | 22 |
| $D$ | Number of acidic groups per mol of GAGs | 1 mol | Estimate |
| $E_f$ | Fiber modulus of collagen network (2.5 mg/mL) | 100 Pa | Fit to experiment |
| $\mu_{col}$ | Shear modulus of collagen network (2.5 mg/mL) | 80 Pa | Fit to experiment |
| $\kappa_{col}$ | Bulk modulus of collagen network (2.5 mg/mL) | 100 Pa | Fit to experiment |
| $m$ | Collagen stiffening exponent | 20 | 2, 3, 18 |
| $n$ | Collagen stiffening exponent | 10 | 2, 3, 18 |
| $\lambda_c$ | Critical stretch for collagen alignment | 1.02 | 2, 3, 18 |
| $\alpha$ | Stiffening exponent of collagen upon swelling | 4.5 | Fit to experiment |

**Figure S3.**
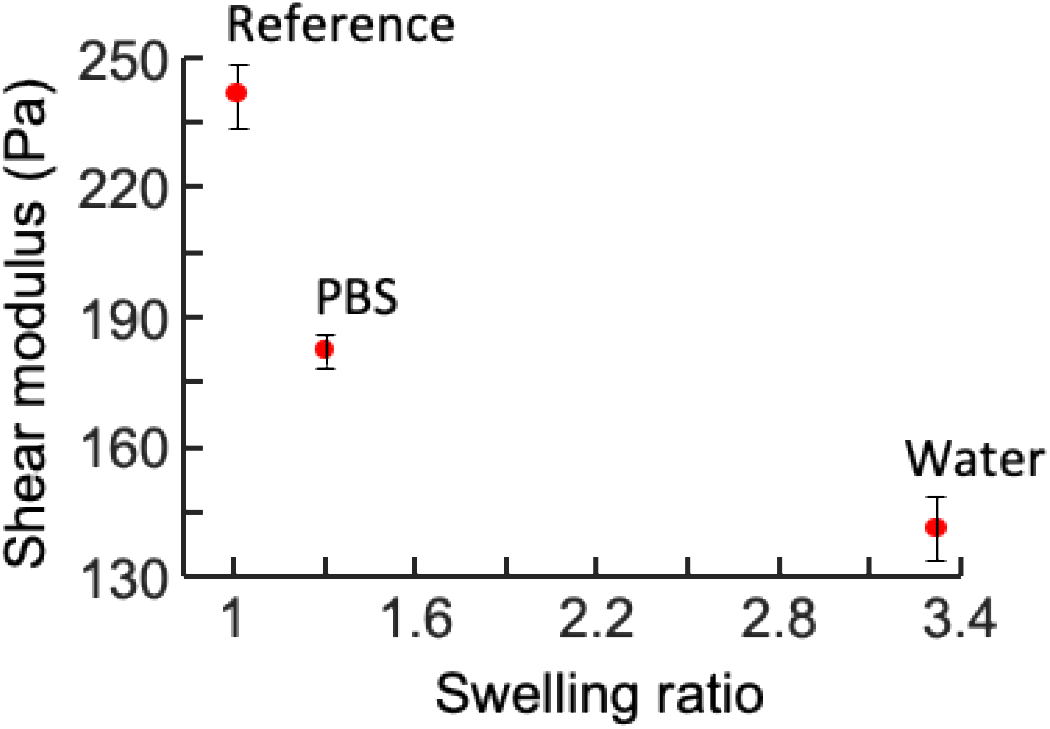
Stiffness of crosslinked HA gel as a function of swelling ratio.

**Figure S4.**
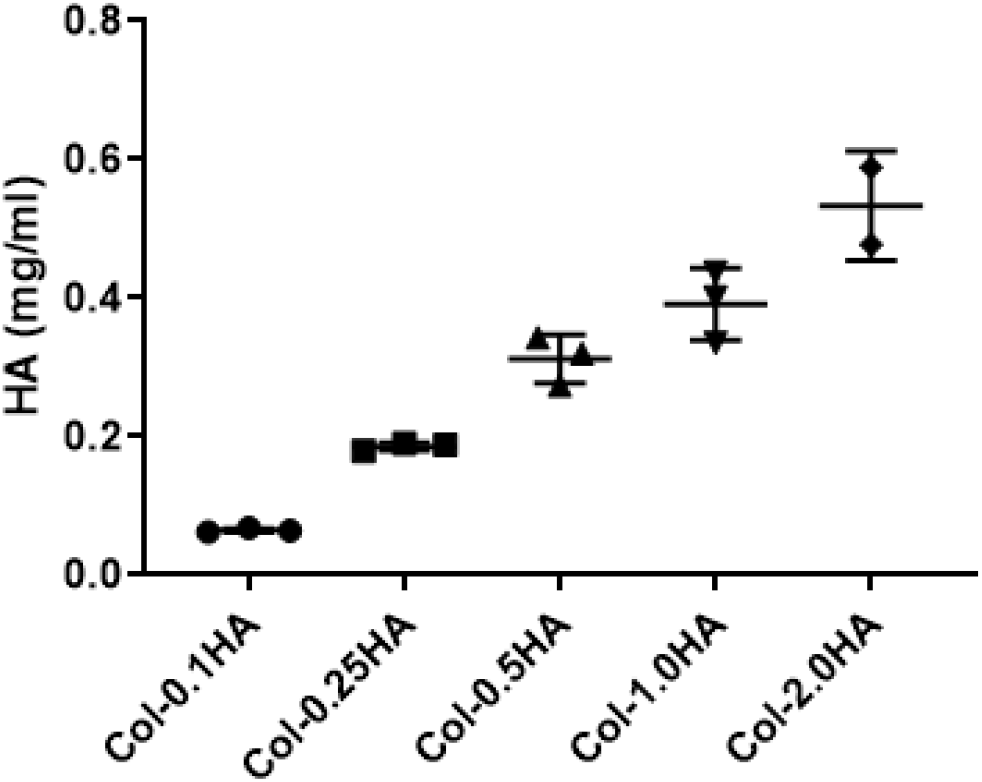
Amount of HA that remained in the collagen-HA co-gel after swelling.

**Figure S5.**
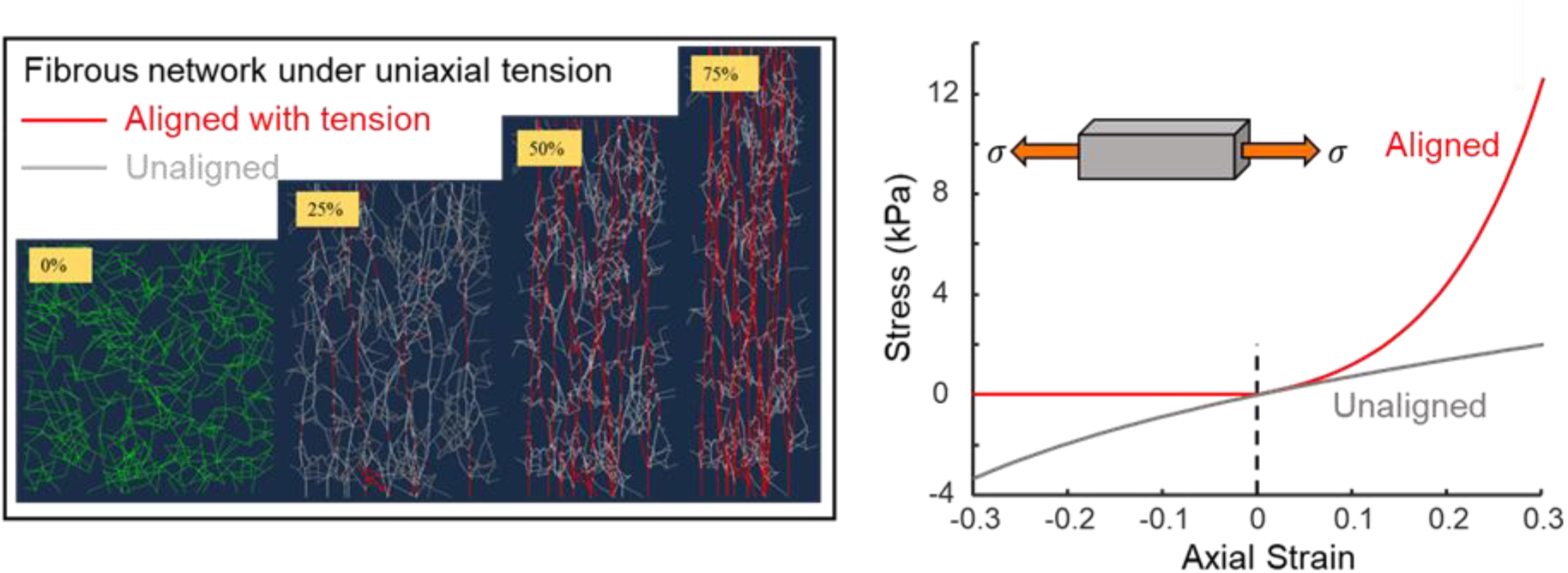
Alignment of the collagen under uniaxial tension (Left). The stress from the aligned fibers and randomly-aligned fibers (Right). The highly non-linear function *f*() captures the stiffening behavior of the collagen network.

